# Vascular Endothelial Barrier Protection Prevents Atrial Fibrillation by Preserving Cardiac Nanostructure

**DOI:** 10.1101/2023.06.20.545742

**Authors:** Louisa Mezache, Andrew Soltisz, Scott R. Johnstone, Brant E. Isakson, Rengasayee Veeraraghavan

**Author notes:** Corresponding Author:Rengasayee Veeraraghavan, Ph.D. Associate Professor Dept. of Biomedical Engineering The Ohio State University 460 Medical Center Dr., Rm 415A, IBMR Columbus, Ohio 43210. Conflicts of Interests: **None.**. **Availability of data and material:** Raw experimental data are backed up on OSU servers and will be shared freely on request. **Code availability:** Image analysis and simulation code will be shared via open forums such as MATLAB Central File Exchange. Twitter handle: @nanocardiology. Tweet: We demonstrate how vascular barrier protection using clinically-relevant drugs can protect cardiac nanostructure and thereby, prevent atrial fibrillation in a setting of acute inflammation.

## Abstract

Atrial fibrillation (AF) is the most common cardiac arrhythmia, affecting ∼3% of the US population. It is widely associated with inflammation, vascular dysfunction, and elevated levels of the vascular leak-inducing cytokine, vascular endothelial growth factor (VEGF). The mechanism underlying AF is not well understood and current treatments are limited to managing this progressive disease, rather than arresting the underlying pathology. We previously identified edema-induced disruption of sodium channel (Na_V_1.5) –rich intercalated disk (ID) nanodomains as a novel mechanism for AF initiation secondary to acute inflammation. Therefore, we hypothesized that protecting the vascular barrier can prevent vascular leak-induced atrial arrhythmias. We identified two molecular targets for vascular barrier protection, connexin43 (Cx43) hemichannels and pannexin-1 (Panx1) channels, which have been implicated in cytokine-induced vascular leak. AF incidence was increased in untreated mice exposed to VEGF relative to vehicle controls. VEGF also increased the average number of AF episodes. VEGF shifted Na_V_1.5 signal to longer distances from Cx43 gap junctions (GJs), measured by a distance transformation-based spatial analysis of 3D confocal images of IDs. Similar effects were observed with Na_V_1.5 localized near mechanical junctions (MJs) composed of N-cad. Blocking connexin43 hemichannels (αCT11 peptide) or Panx1 channels (PxIL2P peptide) significantly reduced the duration of AF episodes compared to VEGF alone with no treatment. Concurrently, both peptide therapies preserved Na_V_1.5 distance from GJs to control levels and reduced MJ-adjacent intermembrane distance in these hearts. Notably, similar antiarrhythmic efficacy was also achieved with clinically-relevant small molecule inhibitors of Cx43 and Panx1.

## INTRODUCTION

Atrial fibrillation (AF) is the most common type of cardiac arrhythmia, affecting 3 to 6 million people in the US^1^. Left untreated, AF progresses from spontaneous episodes to permanent events, and predisposes patients to stroke and more severe cardiovascular disease, impairing quality of life. Currently available treatments primarily entail managing symptoms and often involve more invasive procedures with greater risks and limited efficacy. Interventions such as electrical cardioversion and catheter ablation are expensive, require major clinical infrastructure, and yet, AF recurrence of up to 80% has been reported following these therapies^2, 3^. And though antiarrhythmic drugs used to control rate and rhythm improve symptoms, many have potential deleterious and life-threatening side effects^4^. Interestingly, combining such pharmacological treatments with procedures does not provide a substantial improvement in care nor does it mitigate AF recurrence rate (∼65%)^3^. The critical barrier to progress is that available AF therapies do not target the underlying disease process, and thus are not effective treatment options.

Notably, AF is associated with inflammation, vascular dysfunction, and cardiac structural remodeling^5–7^. Early-stage AF patients have elevated serum levels of proinflammatory cytokines – vascular endothelial growth factor (VEGF), TNFα, and IL6 to name a few – which promote vascular leak and edema^8^. We have previously identified a novel arrhythmia mechanism in which VEGF-induced vascular leak promotes atrial arrhythmias by disrupting sodium channel (Na_V_1.5)-rich intercalated disk (ID) nanodomains^9^. Therefore, we hypothesized that protecting the vascular endothelial barrier can prevent vascular leak-induced atrial arrhythmias. While inflammatory cytokines act through myriad pathways, confounding therapeutic targeting, downstream bottlenecks in the signaling process leading to breakdown of endothelial tight junctions could present attractive alternatives. Connexin 43 hemichannels (Cx43 HCs) and pannexin1 (Panx1) channels play an important role in vascular endothelial barrier integrity, responding to signaling molecules and regulating vascular permeability. During periods of stress, or in regions of injury, these non-specific channels remain open as part of the inflammatory response. Several studies have reported vasculoprotective effects with Cx43 HC and Panx1 channel inhibition in both cardiac^10–13^ and non-cardiac tissues^14, 15^. Thus, we investigated the potential of these inhibitors as antiarrhythmic therapies in our vascular leak-induced atrial arrhythmia model. Here, we provide structural and functional evidence demonstrating that inhibiting these channels exhibits cardioprotective effects.

## METHODS

All animal procedures were approved by Institutional Animal Care and Use Committee at The Ohio State University and performed in accordance with the Guide for the Care and Use of Laboratory Animals published by the U.S. National Institutes of Health (NIH Publication No. 85- 23, revised 2011).

### In vivo ECG

Continuous ECG recordings (PL3504 PowerLab 4/35, ADInstruments) were obtained from mice anesthetized with isoflurane (1-1.5%) as previously described^16^. Briefly, after baseline recording (5 min.), animals received either intraperitoneal VEGF (10 or 50 ng/kg; Sigma) or vehicle (PBS). After an additional 20 min, animals were injected intraperitoneally with epinephrine (1.5 mg/kg; Sigma) and caffeine (120 mg/kg; Sigma) challenge and ECG recording continued for 40 minutes. ECG recordings were analyzed using the LabChart 8 software (ADInstruments). At the conclusion of each ECG recording, the heart was excised, leading to euthanasia by exsanguination. The isolated hearts were prepared in one of the following three ways:

1. <u>Cryopreservation</u>: Hearts were embedded in optimal cutting temperature compound and frozen using liquid nitrogen for cryosectioning and fluorescent immunolabeling as in previous studies^17–20^. These samples were used for light microscopy experiments as described below.
2. <u>Fixation for Transmission Electron Microscopy (TEM)</u>: Isolated hearts were perfused fixed with 2% paraformaldehyde. Atria were then dissected and fixed overnight in 2.5% glutaraldehyde at 4°C for resin embedding and ultramicrotomy as previously described^18, 19^.

### Primary Antibodies

The following primary antibodies were used for Western immunoblotting and fluorescence microscopy studies:

- connexin43 (Cx43; rabbit polyclonal; Sigma C6219)
- connexin43 (Cx43; mouse monoclonal; EMD Millipore Corp. MAB3067)
- N-cadherin (N-cad; mouse monoclonal; BD Biosciences 610920)
- cardiac isoform of the voltage-gated sodium channel (Na_V_1.5; rabbit polyclonal; custom antibody^18^)

### Fluorescent Immunolabeling

Immuno-fluorescent labeling of cryosections (5 µm thickness) of fresh-frozen myocardium was performed, as previously described ^16, 18, 19, 21^. Briefly, cryosections were fixed with paraformaldehyde (2%, 5 minutes at room temperature), permeabilized with Triton X-100 (0.2% in PBS for 15 minutes at room temperature) and treated with blocking agent (1% BSA, 0.1% triton in PBS for 2 hours at room temperature) prior to labeling with primary antibodies (overnight at 4°C). Samples were then washed in PBS (3 x 5 minutes in PBS at room temperature) prior to labeling with secondary antibodies.

For confocal microscopy, samples were then labeled with goat anti-mouse and goat anti-rabbit secondary antibodies conjugated to Alexa 488, Alexa 568 and Alexa 647 were used (1:8000; ThermoFisher Scientific, Grand Island, NY). Samples were then washed in PBS (3 x 5 minutes in PBS at room temperature) and mounted in ProLong Gold (Invitrogen, Rockford, IL). For STochastic Optical Reconstruction Microscopy (STORM), samples were labeled with Alexa 647 and Biotium CF 568 fluorophores. STORM samples were then washed in PBS (3 x 5 minutes in PBS at room temperature) and optically cleared using Scale U2 buffer (48 hours at 4°C) prior to imaging^17, 18, 20^.

### Transmission Electron Microscopy (TEM)

TEM images of the ID, particularly gap junctions (GJs) and mechanical junctions (MJs), were obtained at 60,000x magnification on a FEI Tecnai G2 Spirit electron microscope. Intermembrane distance at various ID sites was quantified using ImageJ (NIH, http://rsbweb.nih.gov/ij/), as previously described^18, 19^.

### Confocal Imaging

Confocal imaging was performed using an A1R-HD laser scanning confocal microscope equipped with four solid-state lasers (405 nm, 488 nm, 560 nm, 640 nm, 30 mW each), a 63x/1.4 numerical aperture oil immersion objective, two GaAsP detectors, and two high sensitivity photomultiplier tube detectors (Nikon, Melville, NY). Individual fluorophores were imaged sequentially with the excitation wavelength switching at the end of each frame. Images were collected as z-stacks with fluorophores images sequentially (line-wise) to achieve optimal spectral separation, followed by 3D deconvolution. Images were analyzed using our distance transformation-based spatial analysis approach, Spatial Pattern Analysis using Closest Events (SPACE)^22, 23^.

### Single Molecule Localization

STORM imaging was performed using a Vutara 352 microscope (Bruker Nano Surfaces, Middleton, WI) equipped with biplane 3D detection, and fast sCMOS imaging achieving 20 nm lateral and 50 nm axial resolution, as previously described ^17, 18, 24, 25^. Individual fluorophore molecules were localized with a precision of 10 nm. The two color channels were precisely registered using localized positions of several TetraSpeck Fluorescent Microspheres (ThermoFisher Scientific, Carlsbad, CA) scattered throughout the field of view, with the procedure being repeated at the start of each imaging session. Protein clustering and spatial organization were quantitatively assessed from single molecule localization data using STORM- RLA, a machine learning-based cluster analysis approach, as previously described^17^.

### Statistical Analysis

Treatments were applied in unblinded fashion for all studies. The Kolmogorov-Smirnov test was applied to compare distributions of measurements, while the Wilcoxon Rank Sum test (for TEM data) and a weighted Student’s T-test (for SPACE outputs) were used for comparisons of central tendency. Fisher’s exact test was used to test differences in nominal data. For multiple comparisons, the Bonferroni correction was applied. A p<0.05 was considered statistically significant. All values are reported as mean ± standard error unless otherwise noted.

## RESULTS

### Protecting vascular endothelial barrier reduces AF susceptibility in VEGF-treated hearts

We have previously shown that pro-inflammatory cytokine-mediated vascular barrier breakdown acutely increased AF inducibility^9^, a finding we recapitulate here (Figure 1). Notably, VEGF significantly increased the incidence and duration of atrial arrhythmias which degenerated into AF (Table 1). It should be noted here that the shorter mean and median durations observed for atrial arrhythmias which degenerated into AF in VEGF-treated hearts vs. vehicle controls reflects the marked increase in the incidence of arrhythmias brought on by VEGF (5.5/animal test vs. 0.33/animal tested in vehicle controls), which included numerous short episodes which rapidly degenerated into AF. Thus, we examined whether inhibiting vascular leak was sufficient to prevent atrial arrhythmias in mice injected with clinically-relevant pathological levels of VEGF, as reported in early-stage AF patients^5, 6, 26–28^. Previous studies identify opening of Cx43 HCs and Panx1 channels in the vascular endothelium as key steps in inflammation-induced vascular leak^9^. Importantly, peptide inhibitors of Cx43 HCs (αCT11^11^) and Panx1 channels (PxIL2P^29^) have been shown to blunt pathological vascular leak in non-cardiac tissues. Adult male mice were pre- treated with peptide inhibitors of Cx43 HCs (αCT11; 100µM) and Panx1 (PxIL2P; 1.6 µM) prior to the acute VEGF insult. Representative *in vivo* ECG traces in Figure 1A illustrate atrial arrhythmia elicited by catecholamine challenge in an untreated mouse exposed to VEGF and maintenance of normal sinus rhythm in mice treated with αCT11 or PxIL2P peptide prior to VEGF insult. The AF phenotype was quantified by assessing inducibility, defined as percent of mice tested under each condition that experienced AF (Figure 1B), and the duration AF episodes (Figure 1C). VEGF alone significantly exacerbated the AF phenotype relative to vehicle controls, inducing AF in 70% of mice tested (14/20 vs 3/10 in vehicle controls; Figure 1B) and significantly increasing the number and duration of AF episodes (Figure 1C, Table 1). Although the decrease in AF incidence or burden (total arrhythmia duration per hour of observation; Supplementary Figure 1) in peptide- treated mice compared to mice exposed to VEGF alone did not reach statistical significance, both peptide treatments markedly reduced the number (6 episodes in 10 αCT11-treated mice and 16 episodes in 10 PxIL2P-treated mice vs. 110 episodes in 20 untreated mice exposed to VEGF; Table 1) and maximum duration (387.8s after αCT11 treatment, 125.5 after PxIL2P treatment vs. 620.5s in untreated mice exposed to VEGF, Table 1) of AF episodes compared to VEGF alone with no treatment. We also observed several brief runs of atrial flutter in all groups including untreated controls. Incidence and burden of atrial flutter showed no significant differences across treatment groups (Supplementary Figure 2). Taken together, these data suggest that vascular barrier protection by Cx43 HC and Panx1 channel inhibition can prevent AF in a setting of acute inflammatory insult.

**Figure 1.**
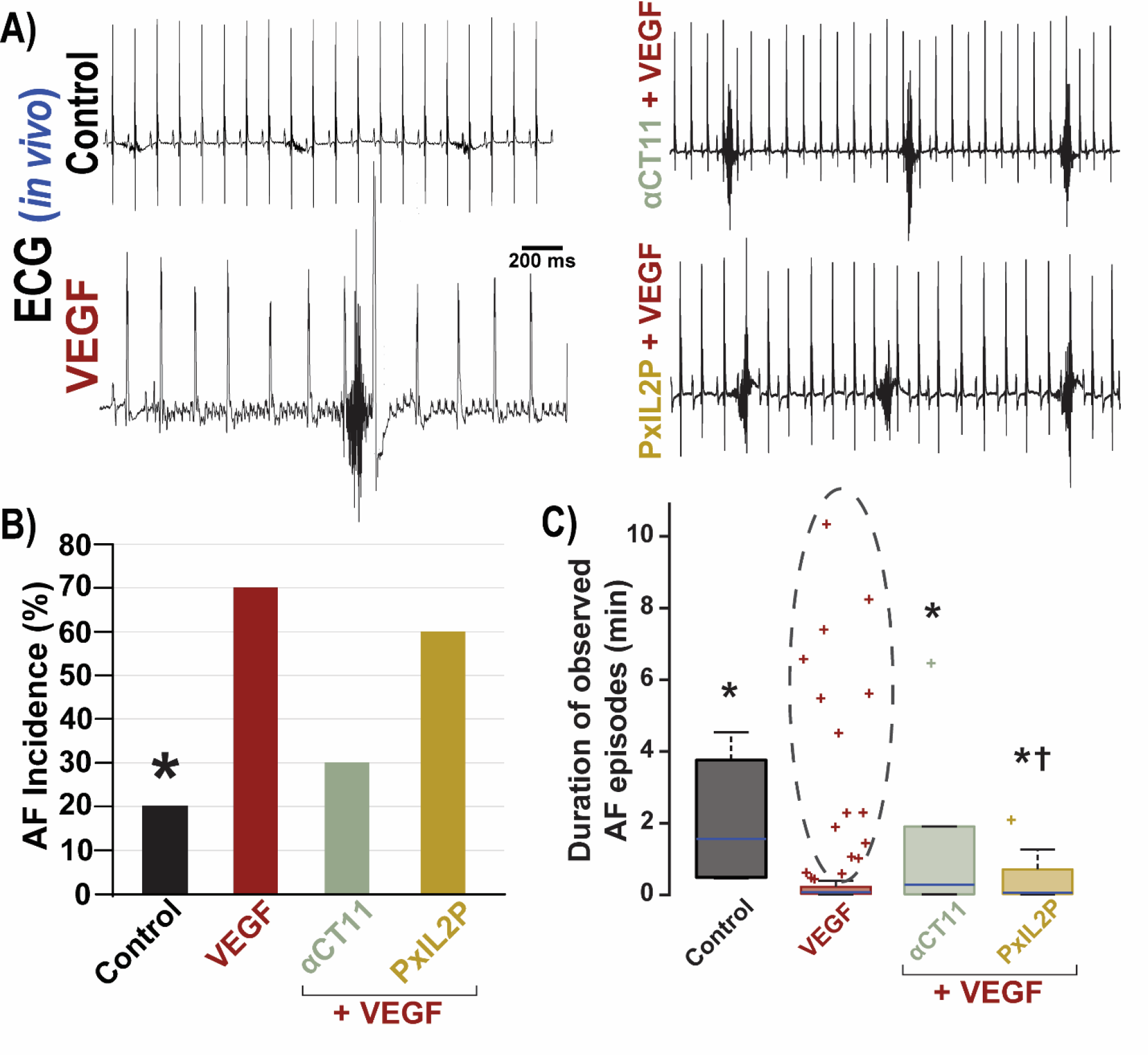
Anti-arrhythmic efficacy of preserving vascular barrier function. A) Representative *in vivo* ECG traces. **B)** Summary plot showing number of atrial fibrillation episodes as percent of mice positive for AF (Control n=15, VEGF n=20, Peptides/Small Molecules n=10 mice/group), * p<0.05 vs. VEGF by Fisher’s Exact test. **C)** Box and whisker plot of AF episode duration, * p<0.05 vs. VEGF by 2 Sample KS test, **_†_** p<0.05 vs. VEGF by Wilcoxon’s test. Dashed black ellipse highlights outliers in the VEGF group, which are longer AF episodes more numerous than all AF episodes observed in other groups. See Table 1 for details.

**TABLE 1.** Characteristics of Atrial Arrhythmias Which Degenerated into AF

| **Characteristics of Atrial Arrhythmias Which Degenerated into AF** |  |  |  |  |  |  |
| --- | --- | --- | --- | --- | --- | --- |
|  | **Control** | **VEGF** | **αCT11** | **PxIL2P** | **Mefloquine** | **Spirolactone** |
| # of experiments | 15 | 20 | 10 | 10 | 10 | 10 |
| # animals that experienced AF | 3 | 14 | 4 | 6 | 1 | 1 |
| # of AF Episodes | 5 | 110 | 6 | 16 | 10 | 8 |
| # of AF Episodes ≥ 30s | 4 | 15 | 2 | 5 | 1 | 0 |
| Average # of episodes/animal tested | 0.33 | 5.50 | 0.60 | 1.60 | 1.00 | 0.80 |
| Average # of episodes/animal experiencing AF | 1.67 | 7.86 | 1.5 | 2.67 | 10.0 | 8.0 |
| Average # of episodes ≥30s/animal experiencing AF | 1.33 | 1.07 | 0.5 | 0.83 | 1 | 0 |
| Mean duration (s) | 126.8 | 37.7 | 89.9 | 23.8 | 8.5 | 4.5 |
| SD (s) | 109.8 | 104.8 | 152.2 | 36.0 | 10.5 | 3.6 |
| p vs. VEGF<br>(Two Sample KS test + Bonferroni's correction) | 1.10E-07 | - | 3.79E-08 | 5.68E-16 | 6.60E-24 | 2.40E-22 |
| Median duration (s) | 93.9 | 4.4 | 17.2 | 3.8 | 2.4 | 4.0 |
| p vs. VEGF<br>(Wilcoxon's test + Bonferroni's correction) | 1.74 | - | 0.415 | 6.62E-07 | 1.96E-15 | 1.95E-13 |
| Max duration (s) | 272.0 | 620.5 | 387.8 | 125.5 | 30.5 | 12.1 |

### Protecting vascular endothelial barrier preserves ID structural integrity following acute VEGF insult

To investigate if the anti-arrhythmic effects of Cx43 HC and Panx1 channel inhibition occur by preventing vascular leak and the ensuing edema-induced ID nanodomain disruption, we performed transmission electron microscopy (TEM) to assess ID structure in mice pretreated with either peptide inhibitor and subsequently subjected to acute VEGF insult. Representative TEM images show intermembrane spacing at GJ- and MJ-adjacent sites for all treatment groups (Figure 2A and B). Overall, acute VEGF insult increased intermembrane spacing at perinexal sites as well as near MJ in highly heterogeneous fashion (Figure 2C, D), consistent with our previous results^9^. Both peptide treatments preserved close membrane apposition at MJ-adjacent, but not perinexal, intermembrane sites after VEGF insult as compared to no treatment (Figure 2C). In keeping with these central tendencies, distributions of intermembrane distances near mechanical junctions, but not at perinexi, were left-shifted in the peptide-treated hearts compared with hearts exposed to VEGF alone (Figure 2D). However, it should be noted that VEGF alone without treatment prompted the emergence of a mode around 100 nm in perinexal intermembrane spacing, which was not present in controls or in peptide-treated hearts.

**Figure 2.**
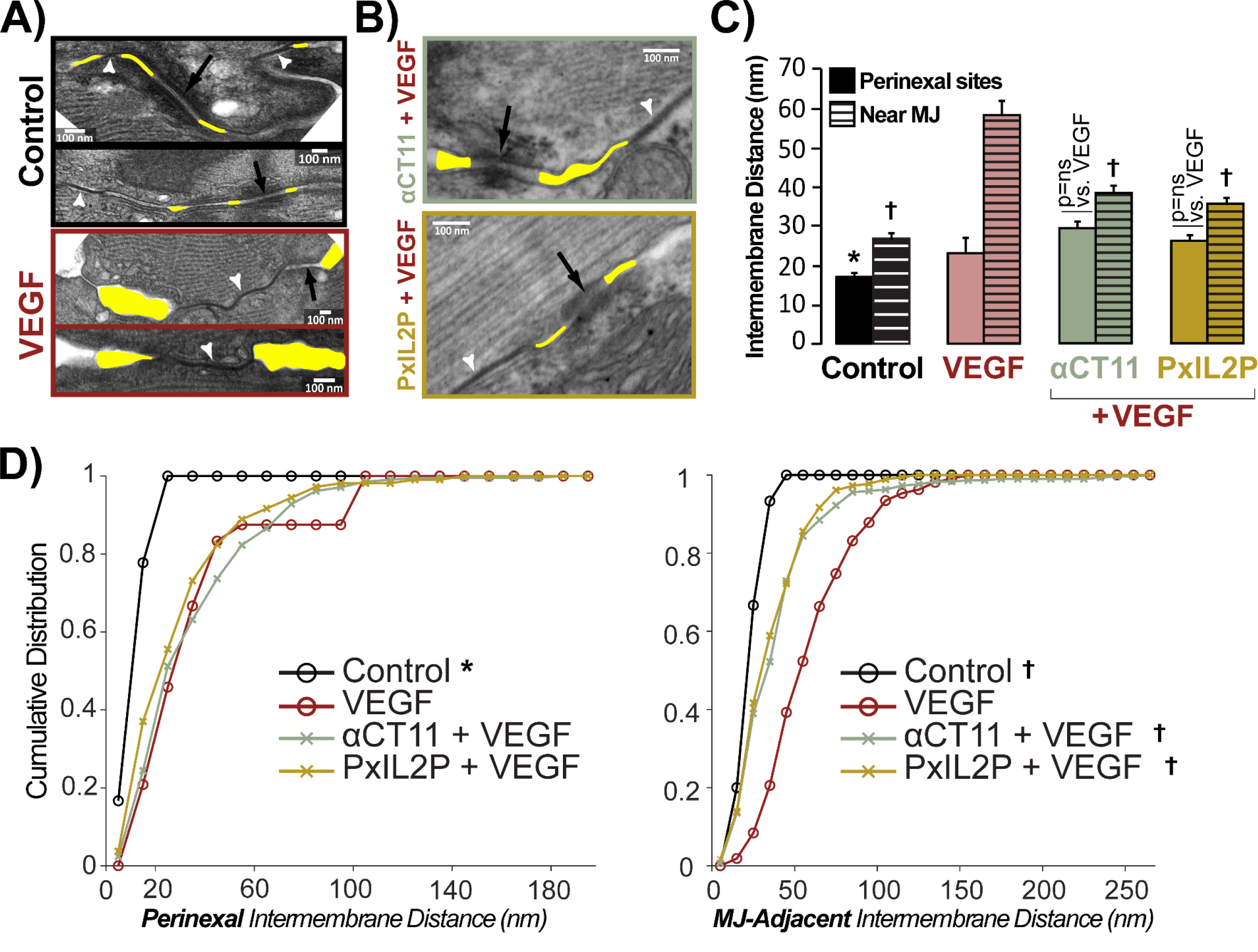
Ultrastructural effects of vascular endothelial barrier protection. Representative TEM images of IDs murine atria. **A)** Untreated control and VEGF. Reproduced from Mezache et al. (2020). **B)** αCT11 and PxIL2P. White arrows point to GJs and black arrows point to MJs. **C)** Summary plots of median intermembrane distance at GJ-adjacent perinexal sites (solid bars) and MJ-adjacent (striped bars) ID sites (>100 measurements/group/location from n=3 hearts/group, * p<0.05 vs. VEGF). **D)** Cumulative distribution of intermembrane distance at perinexal and MJ-adjacent sites. Note the emergence of a second mode at ∼100 nm in perinexal intermembrane distance in the VEGF group, a feature not present in αCT11 / PxIL2P-treated hearts.

### Protecting vascular endothelial barrier preserves Na_V_1.5 localization within the ID following acute VEGF insult

We previously showed Na_V_1.5 migration away from GJs and MJs following acute VEGF treatment^9^; so, we used confocal imaging to examine perinexal and MJ-adjacent Na_V_1.5 distribution in mouse atria pre-treated with either Cx43 HC or Panx1 channel peptide inhibitors. Representative 3D confocal images atrial *en face* IDs illustrate Na_V_1.5 distribution relative to ID landmarks, GJs and MJs, across all treatment groups (Figure 3A and B). Overall, consistent with our previous results obtained using super-resolution microscopy^9^, our distance transformation-based spatial analysis^22, 23^ detected translocation of Na_V_1.5 away from GJs and MJs from diffraction-limited confocal images of VEGF-exposed hearts compared to vehicle controls (Figure 3C, D). Both peptide therapies preserved Na_V_1.5 distance from GJs, but not MJs, to control levels as compared to atria treated with VEGF alone (Figure 3C, D). Furthermore, by comparing the observed distribution of nearest-neighbor distances between immunosignals for co-labeled proteins with the distribution predicted under complete spatial randomness, we obtained measures of non-random attraction / repulsion between the co-labeled proteins. VEGF decreased non-random attraction of Na_V_1.5 to GJs and MJs, while both peptide therapies preserved Na_V_1.5 distance from GJs at control levels (Figure 3C, D).

**Figure 3.**
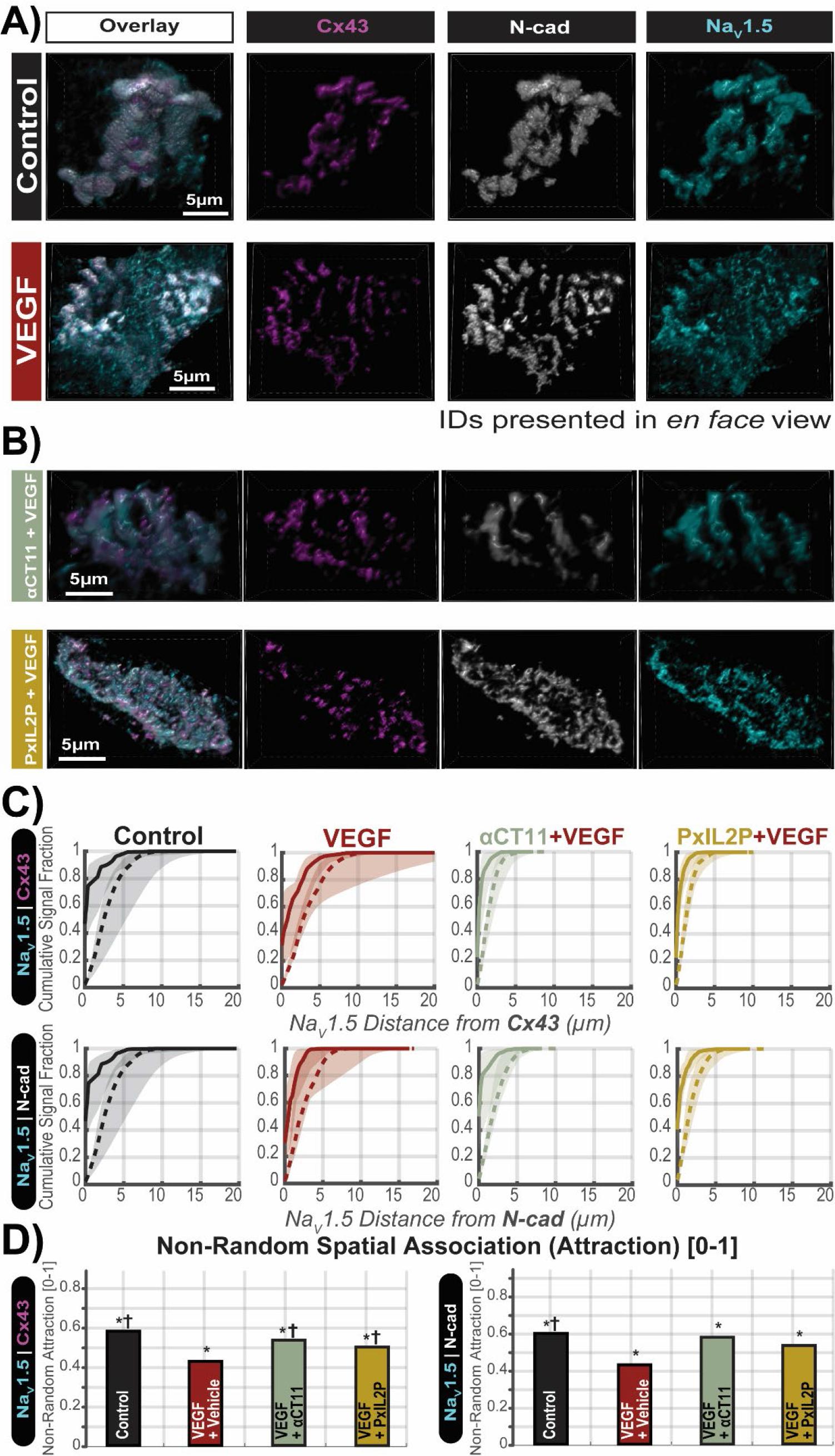
Confocal imaging of IDs. Representative 3D confocal images of *en face* IDs from murine atria. **A)** Untreated controls and VEGF-treated. **B)** Cx43 HC and Panx1 channel peptide inhibitors (αCT11 and PxIL2P, respectively). **C)** Cumulative distribution of Na_V_1.5 distance from Cx43 or Ncad. **D)** Summary plot of Na_V_1.5 attraction to GJs and MJs from n=3 hearts/group, * p<0.05 vs. VEGF by Weighted t-test.

Thus, spatial analysis of 3D confocal images provides measures of Na_V_1.5 organization relative to specific ID landmarks, GJs and MJs. However, under physiological conditions, the distance between clusters of Na_V_1.5 channels and these ID landmarks often falls below Abbe’s diffraction limit. Thus, to directly assess Na_V_1.5 organization within these nanodomains and obtain orthogonal validation of the confocal image results, we turned to STORM single molecule localization microscopy and STORM-RLA machine learning-based cluster analysis. The sub-diffraction resolution of STORM coupled with localization of molecules allows for more discrete measure of Na_V_1.5 protein density at our ID regions of interest, GJs and MJs. Representative 3D *en face* views of atrial IDs show retention of Na_V_1.5 cluster density within 50nm of GJs and MJs in peptide pretreated hearts as compared to untreated controls and VEGF-treated (Figure 4A, B). Overall, both peptide therapies maintained Na_V_1.5 distribution nearer to GJs compared to VEGF (Figure 4C, Supplementary Figures 3, 4). However, there was no significant effect of either peptide on Na_V_1.5 distribution near MJs. These results are consistent with spatial analysis of the diffraction-limited 3D confocal images presented above.

**Figure 4.**
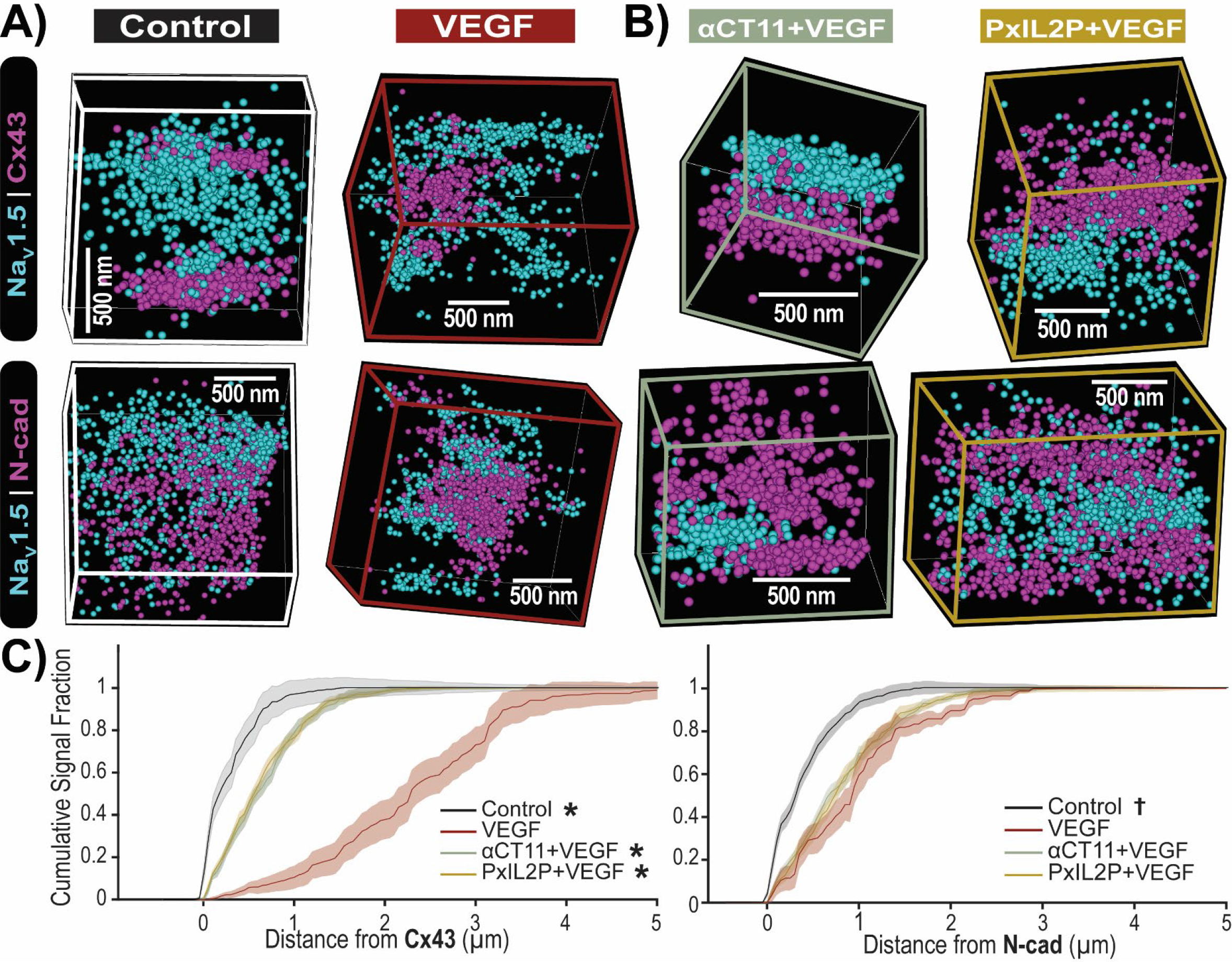
STORM imaging of atrial IDs – Na_V_1.5. Representative 3D STORM images of *en face* IDs immunolabeled for Na_V_1.5 along with Cx43 and N-cad from **A)** Control and VEGF-treated, **B)** peptide pretreated murine atria. STORM data are rendered as point clouds with each localized molecule represented as a 50 nm sphere. Although 20 nm resolution was achieved, the 50 nm size was chosen for rendering to guarantee visibility in print. **C)** Cumulative distribution of Na_V_1.5 expression relative to GJs (Cx43) and MJs (Ncad) from n=3 hearts/group, ^*, †^ p<0.05 vs. VEGF.

### Small molecule alternatives to peptide inhibitors reduce AF susceptibility following acute VEGF insult

While peptide inhibitors offer greater selectivity, *in vivo* delivery of these drugs is poses significant challenges, limiting clinical translatability. Therefore, we identified small molecule alternatives to the Cx43 HC and Panx1 channel peptide inhibitors, mefloquine and spironolactone, respectively. While there are many Cx43 HC and Panx1 channel blockers, mefloquine and spironolactone at the doses used here afford the highest potency with the lowest likelihood of adverse effects, especially gap junction inhibition. Mice were pre-treated with either mefloquine or spironolactone for 10 minutes, followed by acute VEGF insult. Both small molecule treatments significantly reduced AF incidence (Figure 5B, Table 1), episode duration (8.5 ± 10.5s in mefloquine-treated mice, 4.5 ± 3.6 s in spironolactone-treated mice vs. 37.7 ± 104.8s in untreated mice exposed to VEGF; Figure 5C, Table 1), burden (Supplementary Figure 1) and number of episodes (10 episodes in 10 mefloquine-treated mice, 8 episodes in 10 spironolactone-treated mice vs. 110 episodes in 20 untreated mice exposed to VEGF; Table 1). Together, these results suggest that protecting the vascular endothelial barrier reduces atrial arrhythmia susceptibility by preventing edema-induced ID remodeling.

**Figure 5.**
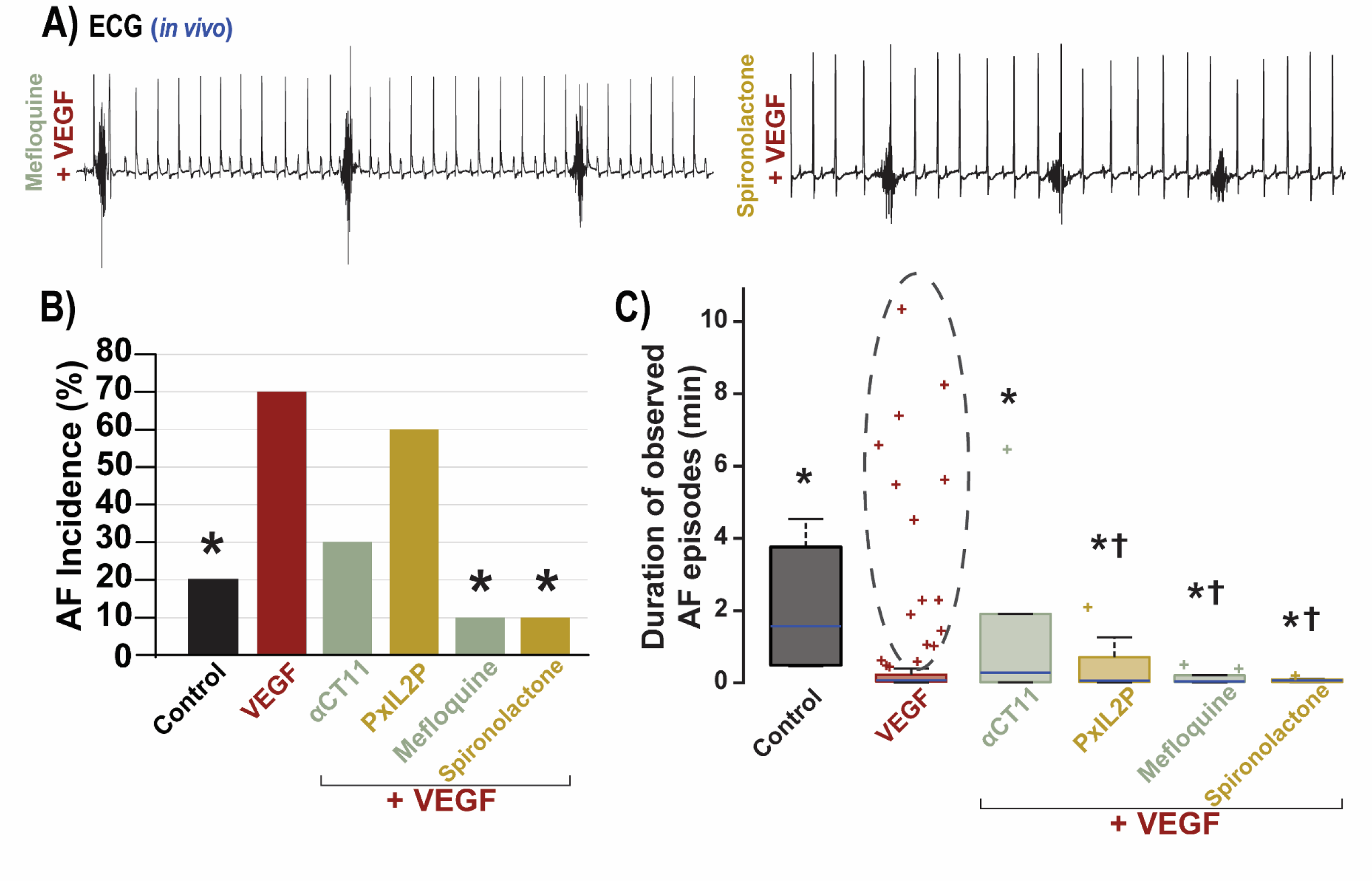
Anti-arrhythmic efficacy of small molecule alternatives to peptide inhibitors. A) Representative *in vivo* ECG traces. **B)** Summary plot showing number of atrial fibrillation episodes as percent of mice positive for AF, * p<0.05 vs. VEGF by Fisher’s Exact test. **C)** Box and whisker plot summarizing duration of AF episodes observed (Control n=15, VEGF n=20, Peptides/Small Molecules n=10 mice/group), * p<0.05 vs. VEGF by 2 Sample KS test, **^†^** p<0.05 vs. VEGF by Wilcoxon’s test. Dashed black ellipse highlights outliers in the VEGF group, which are longer AF episodes more numerous than all AF episodes observed in other groups. See Table 1 for details.

## DISCUSSION

Patients presenting with new onset AF experience inflammation, vascular dysfunction, tissue edema and have acutely elevated levels of vascular leak-inducing cytokines, VEGF^5, 6, 26–28, 30^. We previously identified a novel arrhythmia mechanism by which inflammation-induced vascular leak disrupts Na_V_1.5-rich ID nanodomains, slowing conduction and promoting atrial arrhythmias^9^. Indeed, AF is a common sequela of vascular barrier breakdown in many pathologies. We recognize that inhibiting pro-inflammatory cytokine receptors as an ostensibly evident therapy strategy, however the inflammatory response acts through myriad interconnected pathways, confounding therapeutic targeting. Further, systemic inhibition of inflammatory receptors could interfere with vital physiological processes such as repair after injury or clearing infection. Therefore, we sought to target specific components within endothelial cells, which are critically involved in vascular barrier breakdown, regardless of which inflammatory cytokine may initiate the response. Inflammation-induced vascular barrier breakdown results from dynamic disassembly of tight junctions between vascular endothelial cells in response to inflammation^31^. Key upstream steps involved in this process include the opening of Cx43 HCs^32^ and Panx1 channels^29, 33^ located on the endothelial cell membrane. Under pathological conditions, these large non-specific channels remain open as part of the inflammatory response, activating release of ATP and other small signaling molecules, triggering tight junction breakdown. Notably, endothelial Panx1 has been shown to promote synthesis and release of inflammatory cytokines contributing to positive feedback amplification of inflammatory signaling^34^, further highlighting these channels as a potential therapeutic target. Here, we demonstrate that inhibiting these channels prevents atrial arrhythmias by preserving ID nanodomain structure and protein organization.

Several studies have identified Cx43 HCs and Panx1 channels as crucial contributors to the development of arrhythmogenic substrates^35–39^. And while the mechanism by which these channels increase the risk of cardiac arrhythmias has been largely obscure, inhibiting these channels has demonstrated vasculoprotective and cardioprotective effects. In retina, vascular barrier protection by Cx43 HC and Panx1 channel inhibition prevents retinal damage following ischemic injury and improves vision in patients with age-related macular degeneration and diabetic retinopathy^14, 15^. Inhibition of Cx43 HCs and Panx1 channels has also been shown to reduce inflammation and fibrosis and preserve myocardial function following ischemia-reperfusion injury / myocardial infarction^11, 39–41^. Here, we demonstrate that blocking vascular endothelial Cx43 HCs and Panx1 channels prevents vascular leak, thereby preserving ID nanodomains and reducing arrhythmia susceptibility.

Multiple peptide and small molecule drugs can block Cx43 HCs and Panx1 channels. Cx43 HCs can be inhibited by Gap19^13^ and αCT11^11, 42, 43^, and small molecule drugs such as mefloquine^44^ and tonabersat^14^. While Gap19 has demonstrated protective effects following myocardial ischemia/reperfusion injury, αCT11 inhibits Cx43 HCs by promoting their accretion into GJ plaques, improving coupling between vascular endothelial cells^45^. Furthermore, αCT11 is currently under clinical development to treat dry eye disease as well as ischemia and radiation injury (NCT05031806), facilitating its clinical translation as an antiarrhythmic therapy. Similarly, Panx1 channels can be inhibited by the PxIL2P peptide^29^ and small molecules such as spironolactone^10^ and probenecid^46^. Whereas peptide drugs offer greater selectivity, small molecule inhibitors enable easier *in vivo* delivery. Consistent with other studies, our *in vivo* ECG experiments showed a reduction in atrial arrhythmia burden in mice pre-treated with either Cx43 HC or Panx1 channel peptide inhibitor following acute VEGF insult. Similar antiarrhythmic effects were observed with clinically relevant small molecule alternatives, mefloquine and spironolactone, at doses selected to maximize inhibition while minimizing any off-target effects. Notably, others, who evaluated spironolactone in AF as an aldosterone antagonist, reported antiarrhythmic efficacy in early-stage^12, 47, 48^ but not persistent/permanent^49^ AF patients. This highlights the relevance of a targeted therapeutic approach based on logical, mechanistic understanding of the underlying drivers of AF. Spironolactone was distinctly effective in reducing the risk of new-onset AF, mirroring levels of vascular leak-inducing cytokines, which are only elevated in early-stage AF patients^5, 6, 26–28^, suggesting that it may be uniquely useful in treating early-stage AF. Thus, our results, along with those of the aforementioned clinical trials^12, 47–49^ suggest that spironolactone and mefloquine merit consideration for antiarrhythmic therapy specifically in early stages of AF in a setting of elevated inflammation.

Studies have shown Cx43 HC and Panx1 channel involvement in vascular endothelial permeability^32, 33, 50, 51^, promoting hyperpermeability by the release of key signaling molecules such as ATP in response to inflammatory conditions. As we previously identified in our novel AF mechanism^9^, ID disruption occurs downstream of elevated vascular leak induced by the proinflammatory cytokine VEGF, similar to the ID nanodomain swelling observed in AF patients^7^. Other studies have also demonstrated ID nanodomain swelling following acute interstitial edema^19, 20, 52, 53^. Our TEM studies revealed mitigation of ID nanodomain swelling, particularly MJ- adjacent regions, in hearts pre-treated with either Cx43 HC or Panx1 channel inhibitor following acute VEGF insult. Taken together, these results suggest that preventing inflammation-induced vascular leak preserves ID nanodomain ultrastructure, thereby averting development of a structural proarrhythmic substrate.

Previous studies, including work by our group, have linked the importance of these ID nanodomains in cardiac impulse propagation with the enrichment of Na_V_1.5 sodium channels at GJ-adjacent perinexi and MJ-adjacent sites^17, 19, 20, 54, 55^. Consistent with these reports, we previously demonstrated that VEGF-induced ID disruption and consequent Na_V_1.5 translocation away from these sites was sufficient to induce proarrhythmic conduction slowing^9^. Therefore, we used confocal and super-resolution microscopy to investigate Na_V_1.5 organization at the ID in mice pretreated with Cx43 HC and Panx1 channel inhibitors. Notably, confocal microscopy coupled with spatial analysis^22, 23^ was able to detect VEGF-induced Na_V_1.5 translocation previously identified using super-resolution imaging, specifically, stimulated emission depletion (STED) and STORM single molecule localization microscopy^9^. Consisted with SPACE results from analysis of diffraction-limited confocal images, quantitative analysis of STORM-derived single molecule localizations revealed that our therapeutic peptides preserved measures of Na_V_1.5 attraction to GJs, but not MJs, and Na_V_1.5 cluster density at perinexal sites similar to those of the controls. On a technical level, these results together suggest that spatial analysis of diffraction-limited images could provide clues regarding remodeling occurring at sub-diffraction spatial scales. From a translational perspective, our peptide therapies preserved Na_V_1.5 cluster density near GJs, but did not fully prevent perinexal widening, although they did abolish the emergence of a subpopulation of extremely damaged perinexi with intermembrane distance in excess of 100 nm. Conversely, the peptide therapies maintained normal MJ-adjacent intermembrane distance but not Na_V_1.5 clustering at these sites. Nevertheless, the combined effect of each of the peptide therapies tested was sufficient to ameliorate VEGF-induced atrial arrhythmias in mice.

This points to an important mechanistic implication. Specifically, we demonstrate the antiarrhythmic efficacy of preserving ID nanodomains, which have been shown to play key roles in cardiac action potential propagation by supporting ephaptic coupling: Narrow extracellular cleft width and dense Na_V_1.5 clusters are fundamental requirements for this phenomenon^19, 56, 57^, and previous work by us and others has demonstrated that disruption of these nanodomains results in arrhythmogenic conduction defects^9, 54^. Our electron and light microscopy data suggest that a mitigation of damage to ID nanodomains thought capable of supporting ephaptic coupling is sufficient to protect against arrhythmias. In other words, our therapies did not completely prevent VEGF-induced remodeling of ultrastructure and protein organization within these ID nanodomains, however, they limited heterogeneity to degree compatible with normal atrial rhythm. Collectively, these data suggest protecting the vascular endothelial barrier by inhibiting Cx43 HCs and Panx1 channels prevents ID ultrastructural remodeling and protein reorganization, thereby maintaining functional cardiac ephapses, and preventing the dynamic formation of a structural substrate for arrhythmia (Figure 6).

**Figure 6.**
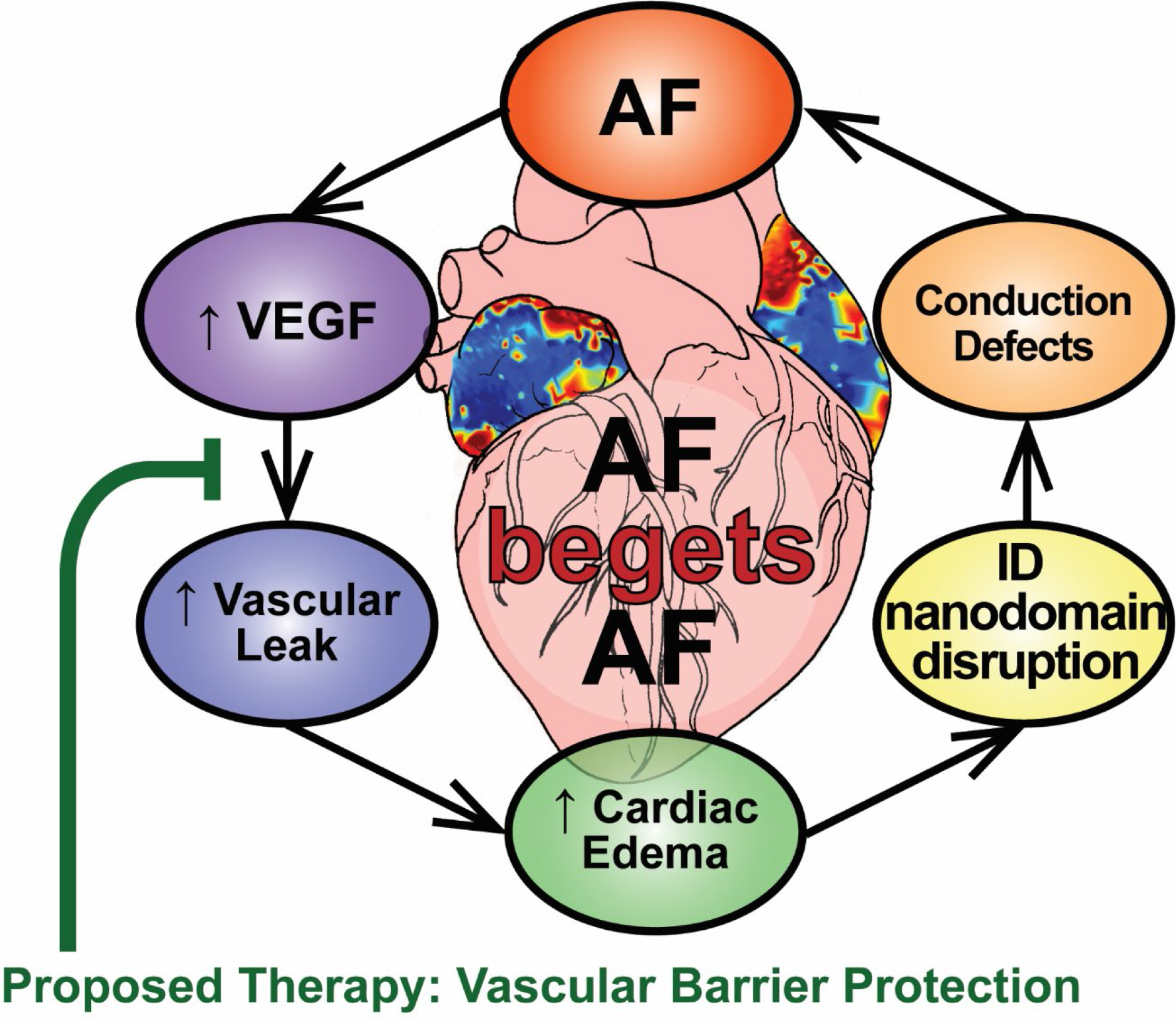
**Proposed strategy for mechanistically derived antiarrhythmic therapy.**

### Limitations

An important concern when targeting Cx43 in the endothelium is the potential impact on Cx43-mediated GJ coupling between cardiomyocytes. However, αCT11 inhibits Cx43 HCs by promoting their accretion into GJ plaques, avoiding any GJ uncoupling^58–61^. While small molecule drugs are easier to deliver *in vivo*, their reduced specificity often yields off-target effects. Among small molecule inhibitors of Cx43 HCs and Panx1 channels, however, mefloquine and spironolactone, respectively present the greatest potency with minimal off-target effects^10, 13, 14, 42, 44, 46, 62–71^. Particularly in the case of Cx43 HC inhibition, the selected dose of mefloquine has no impact on GJ coupling. Interestingly, some studies have demonstrated inhibitory effects of mefloquine on Panx1 channels^72–74^. Based on our ECG studies, the antiarrhythmic effects of mefloquine suggest combined targeting of Cx43 HCs and Panx1 channels is synergistic, though further investigation is needed to fully understand drug interactions.

### CONCLUSIONS

We demonstrate vascular endothelial barrier protection as a mechanistically-derived antiarrhythmic strategy for the prevention of AF in its early stages. This approach, for which we demonstrate both efficacy and mode of action, provides a logical treatment option for arrhythmia prevention in patients at risk for new onset arrhythmias, such as post-op AF. This approach has considerable implications for standard of care for early-stage AF patients, particularly those with underlying pathologies associated with vascular dysfunction.

## CLINICAL PERSPECTIVES

### Competency in Medical Knowledge

Our novel results demonstrate a mechanistically-guided therapeutic approach for the prevention of inflammation-induced atrial fibrillation by protecting the vascular endothelial barrier and thereby, preventing damage to intercalated disc nanodomains vital for cell-to-cell communication.

### Translational Outlook

We provide mechanistic evidence to demonstrate that novel therapeutic strategies, entailing inhibition of connexin 43 hemichannels and pannexin 1 channels in the vascular endothelium using clinically-relevant drugs, can effectively prevent atrial fibrillation in a setting of acute inflammation. This work holds important implications for the prevention of post-operative atrial fibrillation as well as new-onset / early-stage atrial fibrillation associated with inflammatory etiologies.

## ACKNOWLEDGEMENTS

The authors wish to thank Prof. Robert Gourdie for extremely valuable discussions on the biology of connexin 43 hemichannels and the use of mimetic peptides to inhibit them.

## List of Abbreviations

AF: Atrial fibrillation
VEGF: Vascular endothelial growth factor
ID: Intercalated disc
Cx43: Connexin 43
Panx1: Pannexin 1
GJ: Gap junction
MJ: Mechanical junction
NaV1.5: Cardiac isoform of the voltage-gated sodium channel
STORM: Stochastic optical reconstruction microscopy
SPACE: Spatial Pattern Analysis using Closest Events, a spatial image analysis approach

---

**Supplementary Figure 1.**
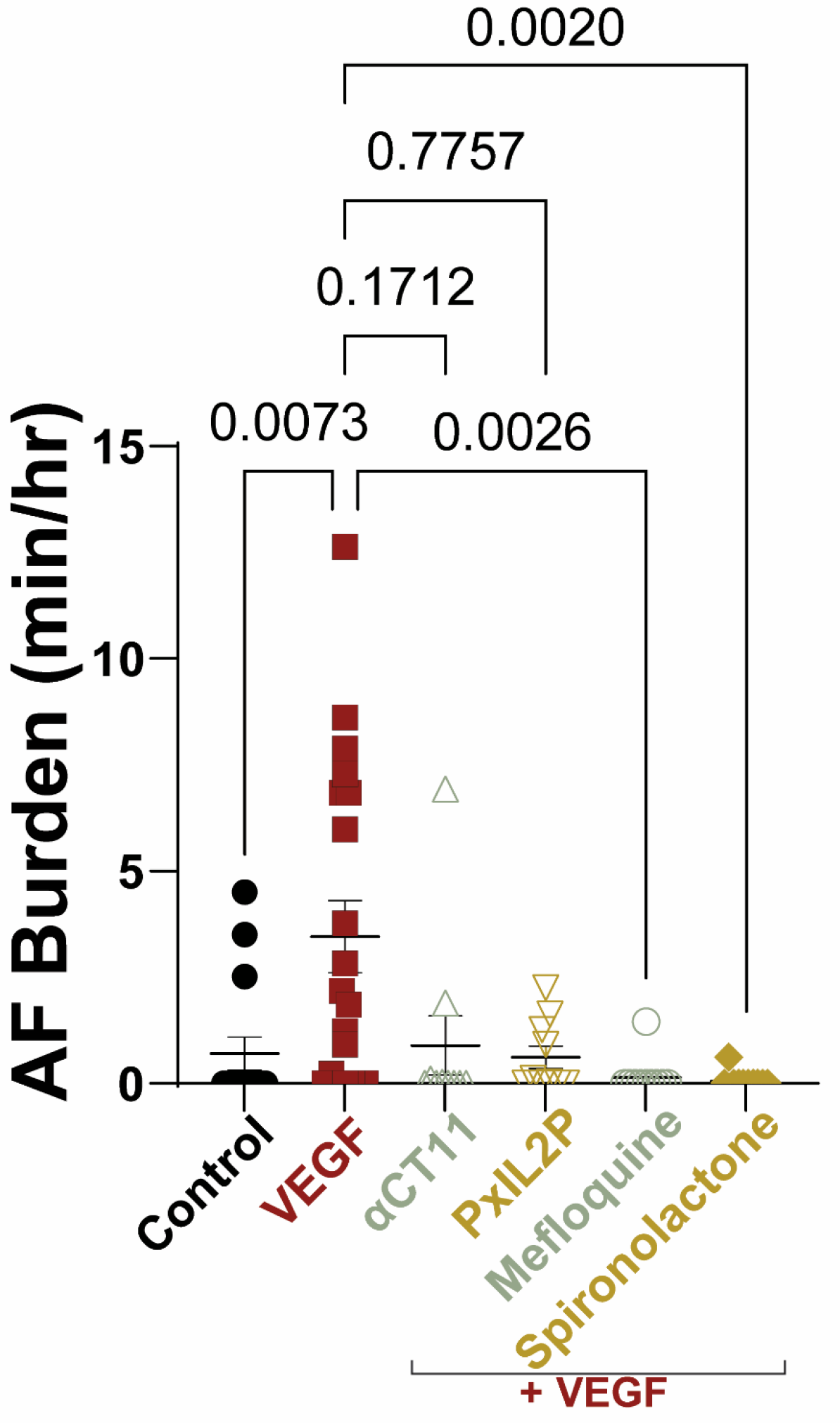
Atrial Fibrillation Burden. A) Total atrial fibrillation burden under caffeine + epinephrine challenge quantified as minutes of fibrillation per hour of observation.

**Supplementary Figure 2.**
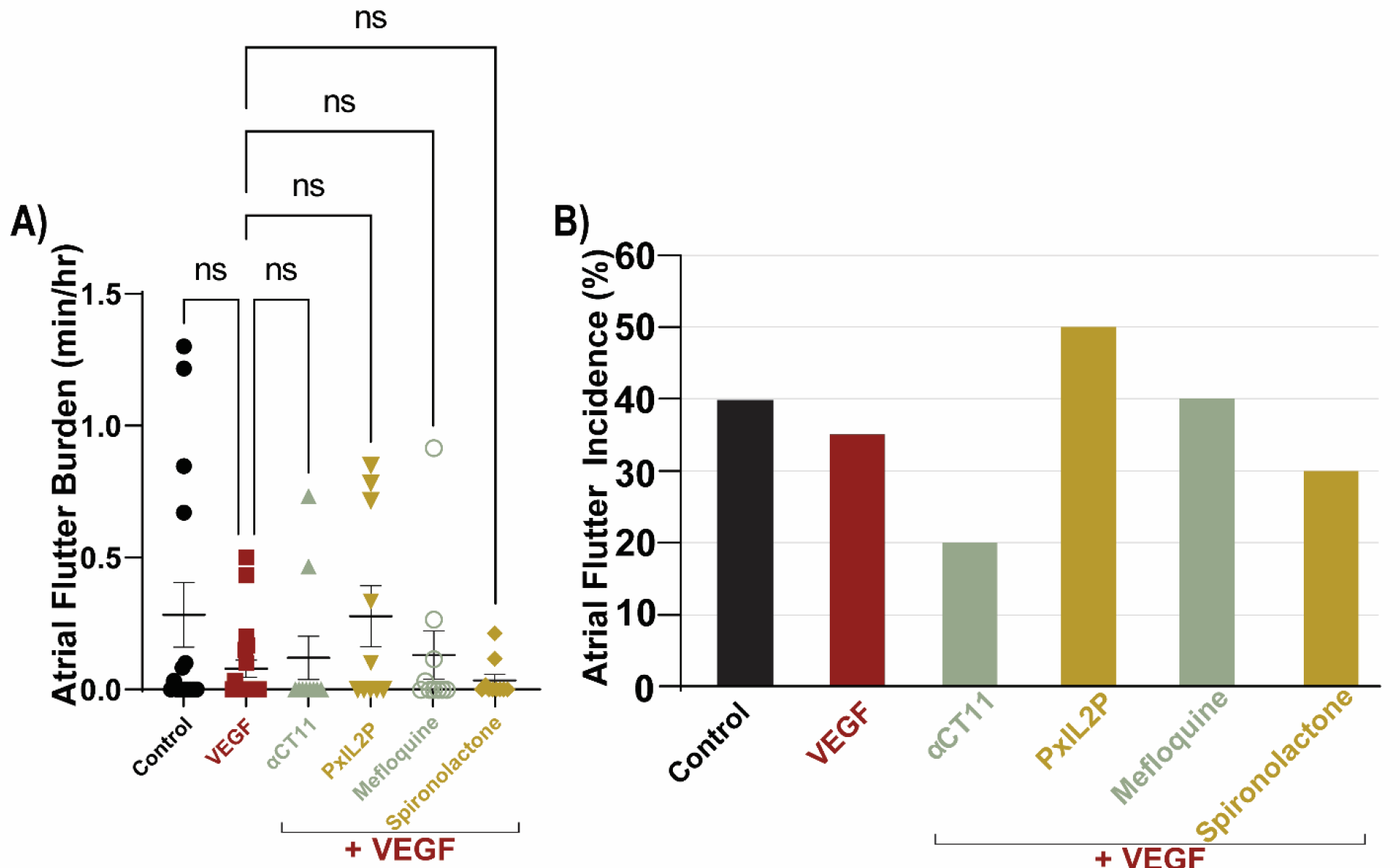
Atrial Flutter Burden and Incidence. A) Total atrial flutter burden under caffeine + epinephrine challenge quantified as minutes of flutter per hour of observation. **B)** Summary plot showing percent incidence of atrial flutter.

**Supplementary Figure 3.**
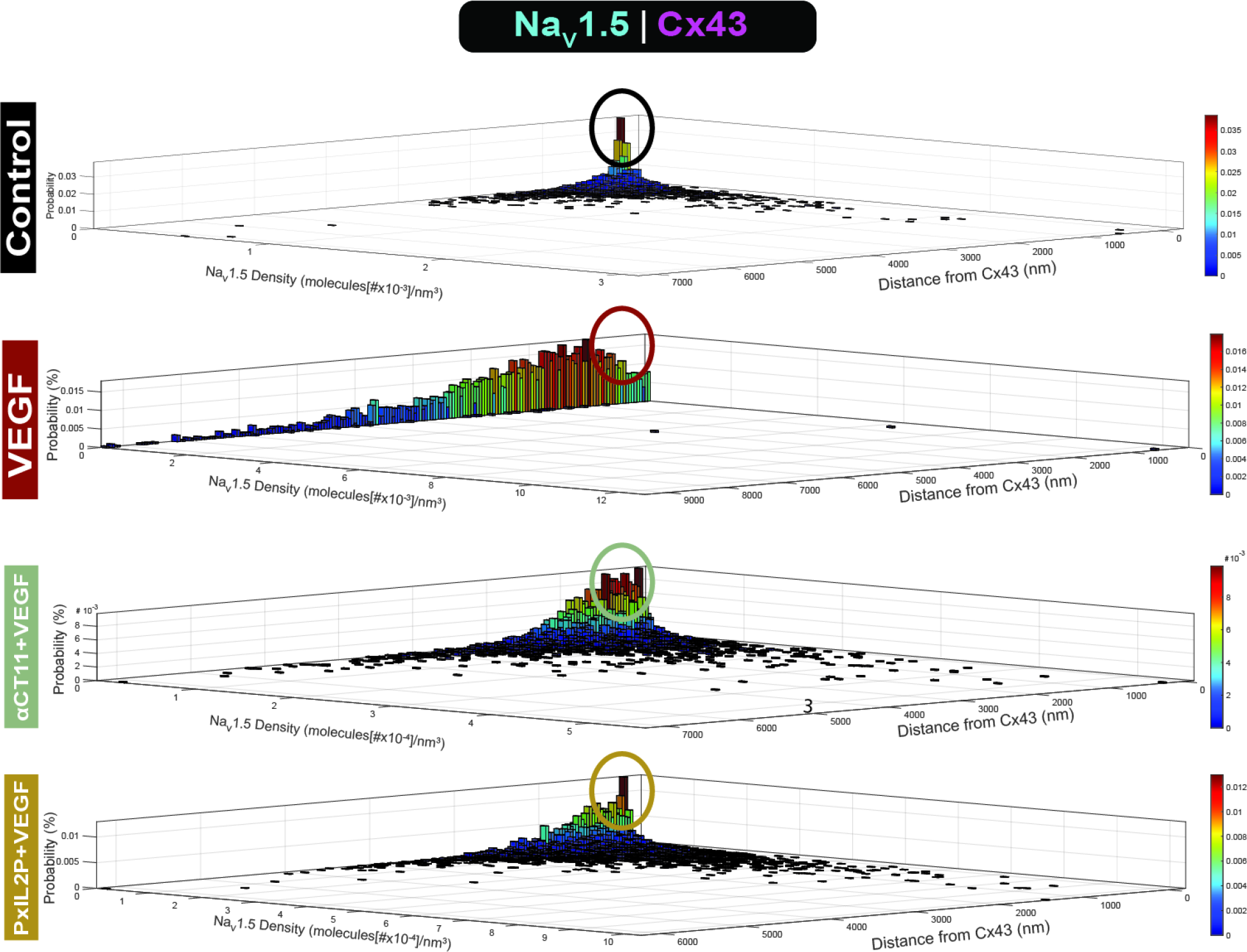
Effects of peptide therapies on perinexal Na_V_1.5 distribution. Summary data demonstrate loss of Na_V_1.5 cluster density within the perinexus in VEGF compared to control (circled region). Na_V_1.5 cluster density at the perinexus is preserved with both peptide therapies (circled region).

**Supplementary Figure 4.**
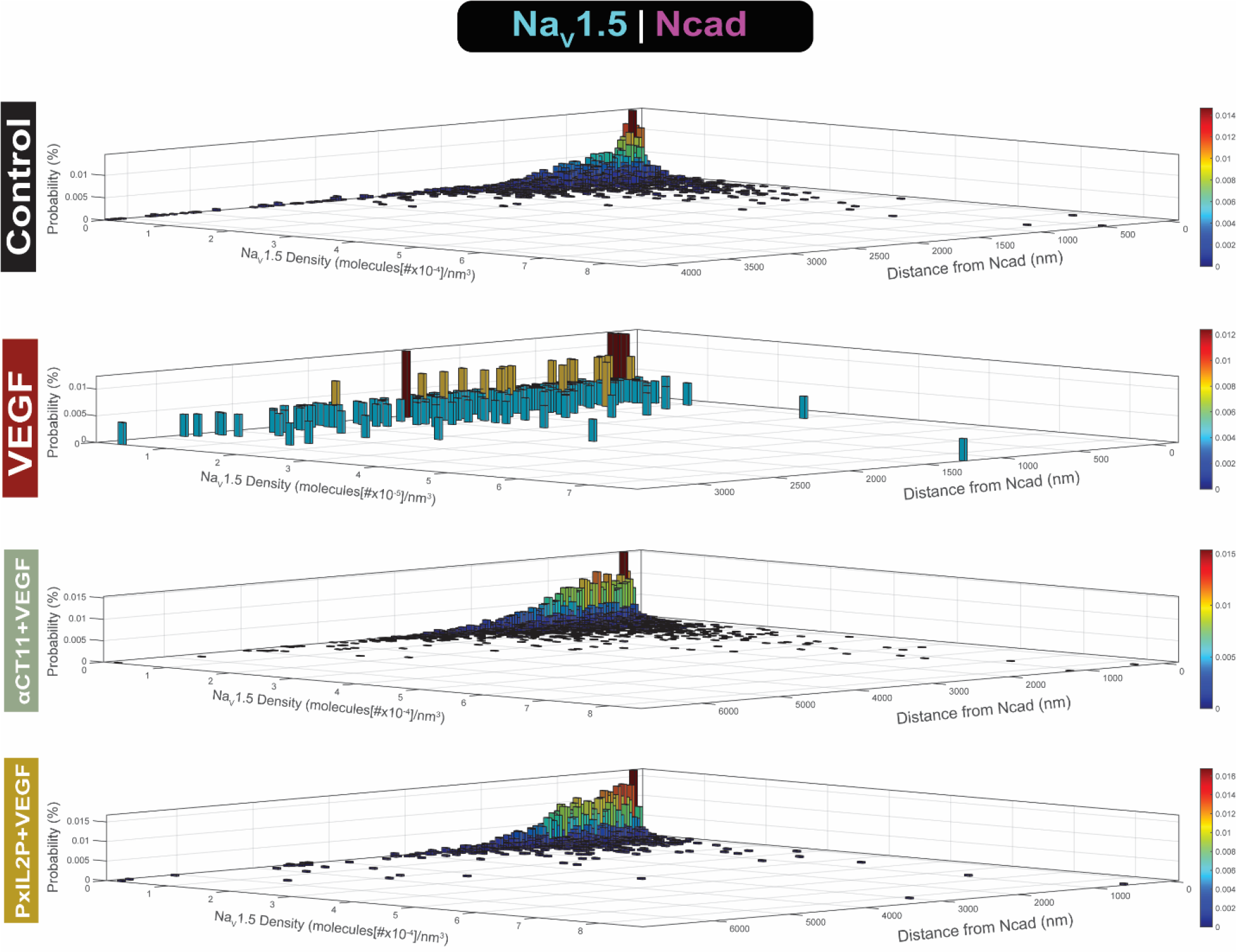
Effects of peptide therapies on MJ-adjacent Na_V_1.5 distribution. Summary data demonstrate increase of Na_V_1.5 expression at greater distances from MJs in VEGF compared to control. Na_V_1.5 cluster density at MJ-adjacent sites is preserved with both peptide therapies.

